# Indication of spatially random infection of chlamydia-like organisms in *Bufo bufo* tadpoles from ponds located in the Geneva metropolitan area

**DOI:** 10.1101/402487

**Authors:** Elia Vajana, Ivo Widmer, Estelle Rochat, Solange Duruz, Oliver Selmoni, Séverine Vuilleumier, Sébastien Aeby, Gilbert Greub, Stéphane Joost

## Abstract

Occurrence of bacteria belonging to the order *Chlamydiales* was investigated for the first time in common toad (*Bufo bufo*) tadpole populations collected from 41 ponds in the Geneva metropolitan area, Switzerland. A *Chlamydiales*-specific Real-Time PCR was used to detect and amplify the *Chlamydiales* 16S rRNA-encoding gene from the tails of 375 tadpoles. We found the studied amphibian populations to be infected by “Chlamydia-like organisms” (CLOs) attributable to the genera *Similichlamydia*, *Neochlamydia*, *Protochlamydia* and *Parachlamydia* (belonging to the family *Parachlamydiaceae*), *Simkania* (family *Simkaniaceae*) and *Estrella* (family *Criblamydiaceae*); additionally, DNA from the genus *Thermoanaerobacter* (family *Thermoanaerobacteriaceae*) was detected. A global autocorrelation analysis did not reveal a spatial structure in the observed CLOs infection rates, and association tests involving land cover characteristics did not evidence any clear effect on CLOs infection rates in *B. bufo*. Despite preliminary, these results suggest a random and ubiquitous distribution of CLOs in the environment, which would support the biogeographical expectation “everything is everywhere” for the concerned microorganisms and their amoeba vectors.

## Introduction

The order *Chlamydiales* consists of strict intracellular bacteria that replicate within eukaryotic cells of several animal hosts, among which humans [1]. Current molecular evidence suggests the existence of two main lineages within *Chlamydiales*, which possibly diverged between 700 and 1400 million years ago from the last common ancestor [2]: the family *Chlamydiaceae*, and the “chlamydialike organisms” (CLOs) belonging to the families *Piscichlamydiaceae*, *Clavichlamydiaceae*, *Simkaniaceae*, *Rhabdochlamydiaceae*, *Waddliaceae*, *Parachlamydiaceae*, *Criblamydiaceae*, and *Parilichlamydiaceae* [1,3,4].

*Chlamydiaceae* were first described in the sixties, and since then they were discovered to cause a wide variety of diseases affecting over 400 documented animal species [1,5,6]. Among the most eminent representatives of the family are (i) *Chlamydia trachomatis*, the etiological agent of the human visually-impairing trachoma [1], (ii) *C. psittaci*, responsible for pneumonia and hepatitis in birds [6], as well as zoonotic lung infections in humans [7] and equine infections [8], (iii) *C. abortus* causing abortion in sheep, goats, cattle and swine, and with proved transferability to humans [6,9], and (iv) *C. pneumoniae*, responsible for respiratory infections and atherosclerosis in humans, rhinitis in koalas and horses, and conjunctivitis in reptiles [1,3,6]. Notably, *C. pneumoniae* was also reported to infect amphibian populations of African clawed frogs (*Xenopus tropicalis*), great barred frogs (*Mixophyes iteratus*), blue mountains tree frogs (*Litoria citropa*), and common frogs (*Rana temporaria*), which were also positive for *C. abortus* and *C. suis* [10–12]; furthermore, the Candidatus *Amphibiichlamydia ranarum* was found at high prevalence in invasive bullfrog populations, and is considered an emerging pathogen possibly contributing to the current amphibian biodiversity crisis [13].

CLOs discovery is more recent, and dates back to the end of the eighties, when *Waddlia chondrophila* was first isolated from an aborted bovine foetus [1,3]. As exhaustively reviewed [3,6,14], most research has focused on identifying emerging pathogens among CLOs, with *W. chondrophila* being suspected to trigger human miscarriage [15] and ruminant abortion [16,17], *Parachlamydia acanthamoebae* human miscarriage [18], *Simkaniaceae*, *Parachlamydiaceae* and *Rhabdochlamydiaceae* respiratory diseases in humans and cattle [19–22], ocular infections in cats [23] and granulomatous inflammation in reptiles [24], *Parachlamydia* species to concur in the massive mortality events affecting an highly endangered midwife toad (*Alytes obstetricans*) population [25], and bacteria belonging to *Clavichlamydiaceae*, *Parachlamydiaceae*, *Parilichlamydiaceae*, *Piscichlamydiaceae*, *Rhabdochlamydiaceae*, and *Simkaniaceae* having a recognized role in epitheliocystis, a common gill disease in fish [3,26,27].

CLOs are also misnamed “environmental chlamydiae”, as the majority of them were first isolated from heterogeneous environmental sources spanning from water to soil [3,28,29]. Notably, CLOs belonging to *Parachlamydiaceae*, *Simkaniaceae*, *Criblamydiaceae* and *Waddliaceae* have been repeatedly observed as obligate endosymbionts of free-living amoebae species of the genera *Acanthamoeba* and *Hartmanella* [5,10,29–31], which are therefore expected to play a central role in guaranteeing CLOs survival and dispersal in the environment, as well as infections in new hosts [32]. Possibly reflecting the high ecological tolerance and dispersal capabilities of such vectors (which can eventually rely on insects and wind to disperse over long distances), CLOs have been isolated from several ecosystems, and are commonly considered ubiquitous in the environment [1,3,26,33,34], with a highly diversified set of hosts including humans, marsupials and small mammals like the fruit bat, reptiles like chelonians, lizards and snakes, fish species like the Leafy sea dragon, the Blue-striped snapper, the Atlantic salmon, and the African catfish, as well as crustaceans like the Rough woodlouse [1,3,9,24,26].

However, biogeographical end ecological studies have been conducted to elucidate possible links between environmental conditions and composition of protozoa communities and distributions, leading to reject the paradigm “everything is everywhere, but, the environment selects” associated with free-living protozoa in some species (see the cases of *Nebela vas* and *Badhamia melanospora*) [35,36]. Particularly, local trends in precipitation [37] and soil characteristics related to moisture, temperature, pH, dissolved oxygen, and land cover (especially in terms of bryophyte species occurring in the crust) showed association with some testate [38–40] and protosteloid amoebae [37] occurrence. Such evidences would suggest – or at least do not exclude – a possible and still largely unexplored environmental influence on CLOs vectors and their endosymbionts at a local geographical scale.

In the present work, we investigated the occurrence of CLOs infection in the widespread common toad (*Bufo bufo*) for the first time, and tested the “everything is everywhere, but, the environment selects” principle with the observed infection patterns [41,42]. Particularly, we first tested *B. bufo* tadpole populations from the Geneva metropolitan area (Switzerland) for infection, and then the resulting infection rates for random distribution and association with land cover characteristics. In a public health perspective, we also derived human population density around sampling sites and studied a possible relationship with *B. bufo* infection, given the CLOs ability to infect humans from environment [3,21,43].

## Materials and Methods

### Sampling

In the context of the URBANGENE project, sampling locations were chosen in the state of Geneva on the basis of the MARVILLE (http://campus.hesge.ch/mareurbaine/) and of the *Centre de coordination pour la protection des amphibiens et des reptiles de Suisse* (http://www.karch.ch/) ponds databases, and also by means of a crowdsourcing campaign to include private ones (http://urbangene.heig-vd.ch). One hundred and fifty ponds were identified and then inspected. Tadpoles were finally sampled from April 9 to 22, 2015, in a subset of 41 ponds (Figure 1). Sampled ponds differed in size, species composition, and in the typology of the surrounding environments, some of them being located in close proximity to the densely inhabited Geneva downtown (and placed in urban parks and private grounds), some others in the more rural Geneva suburbs, characterized by a higher degree of naturalness. Overall, 375 tadpoles were sampled, with an average of 9.2 samples per pond (sampling range: 4-15 tadpoles per pond). In order to characterize the whole tadpole population present in a pond, sampling privileged tadpoles coming from different frogspawns, whenever present; in such a case, tadpoles were collected shortly after they hatched from their frogspawn to reduce the chance of sampling siblings.

**Figure 1.**
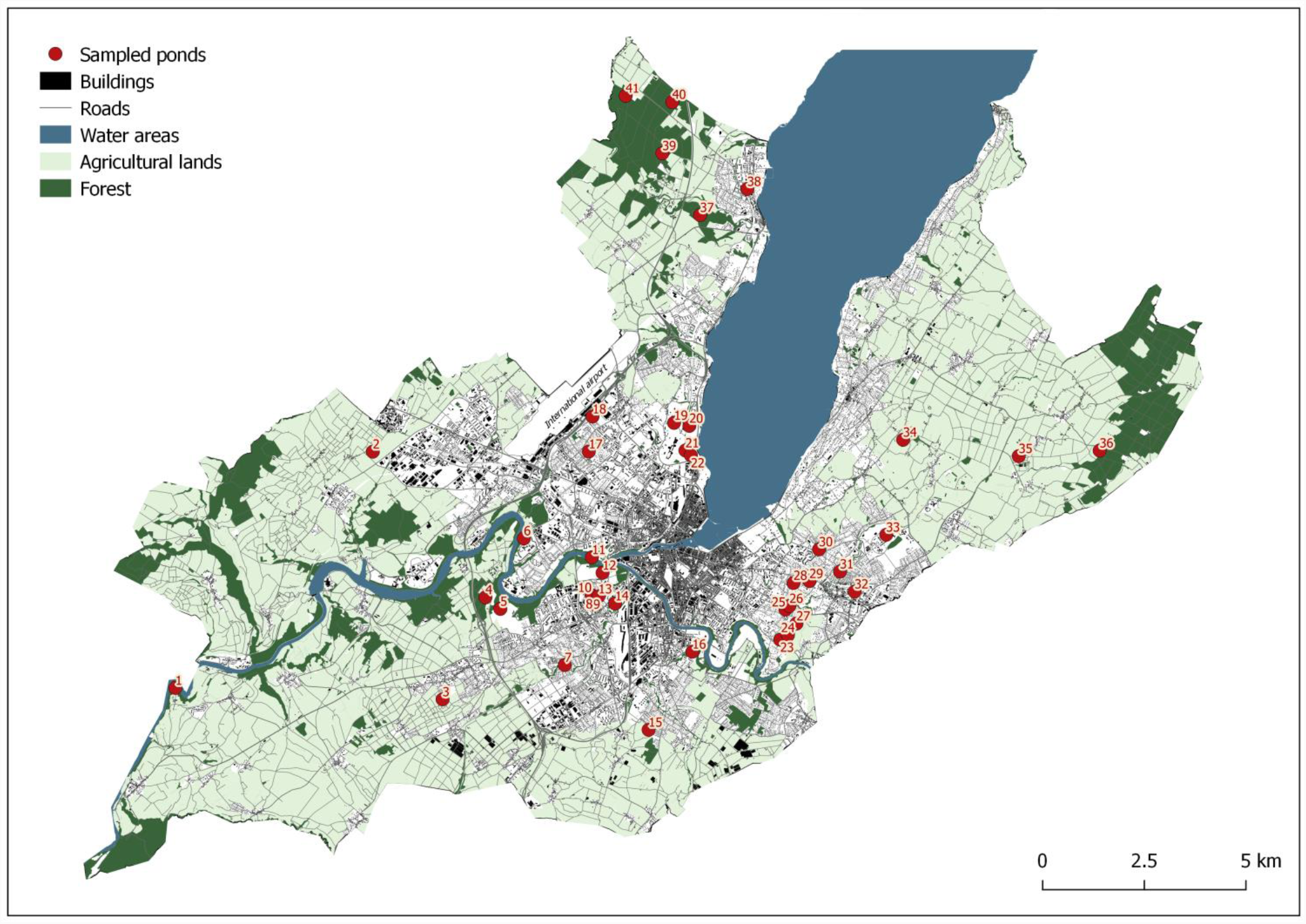
Spatial representation of the sampled ponds. Ponds are represented in red, with numbers corresponding to Pond IDs in Table 1. Background contextual information represents Geneva metropolitan area, and highlights forests, agricultural, water, as well as urbanized areas.

### DNA extraction

#### Sample preparation

After sampling, tadpoles were put individually in a water dish with Tricaine methane sulphonate (MS-222), which caused tadpoles’ rapid anaesthesia and decease. The apical part of the tail was carefully clipped in order to avoid contamination by bacteria from the intestinal tract. After the freeze-drying of the tails, DNA was extracted at the LGC laboratories in Berlin (Germany), using the sbeadex™ tissue kit (LGC, Teddington, UK), and following the manufacturer’s instructions.

**Table 1.**
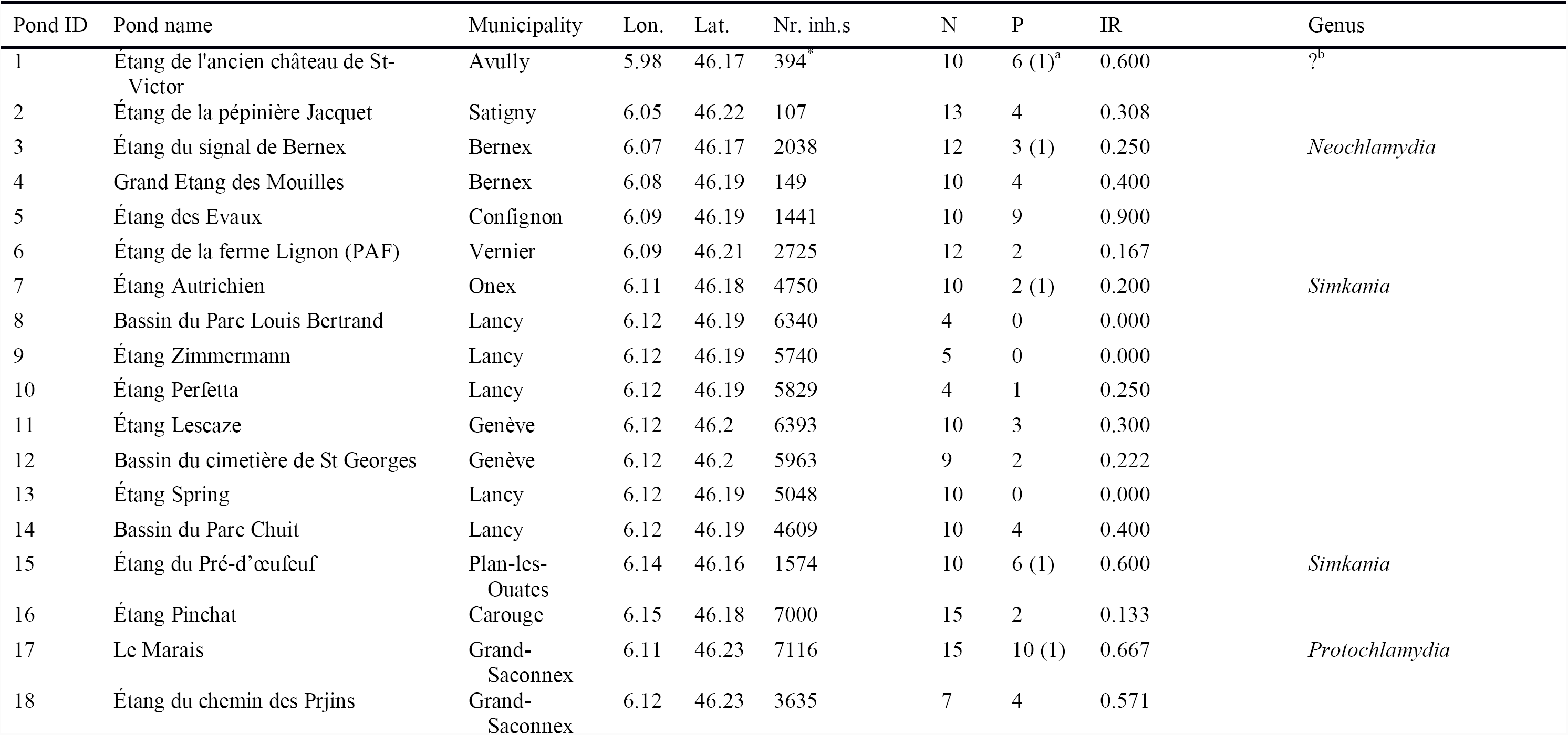
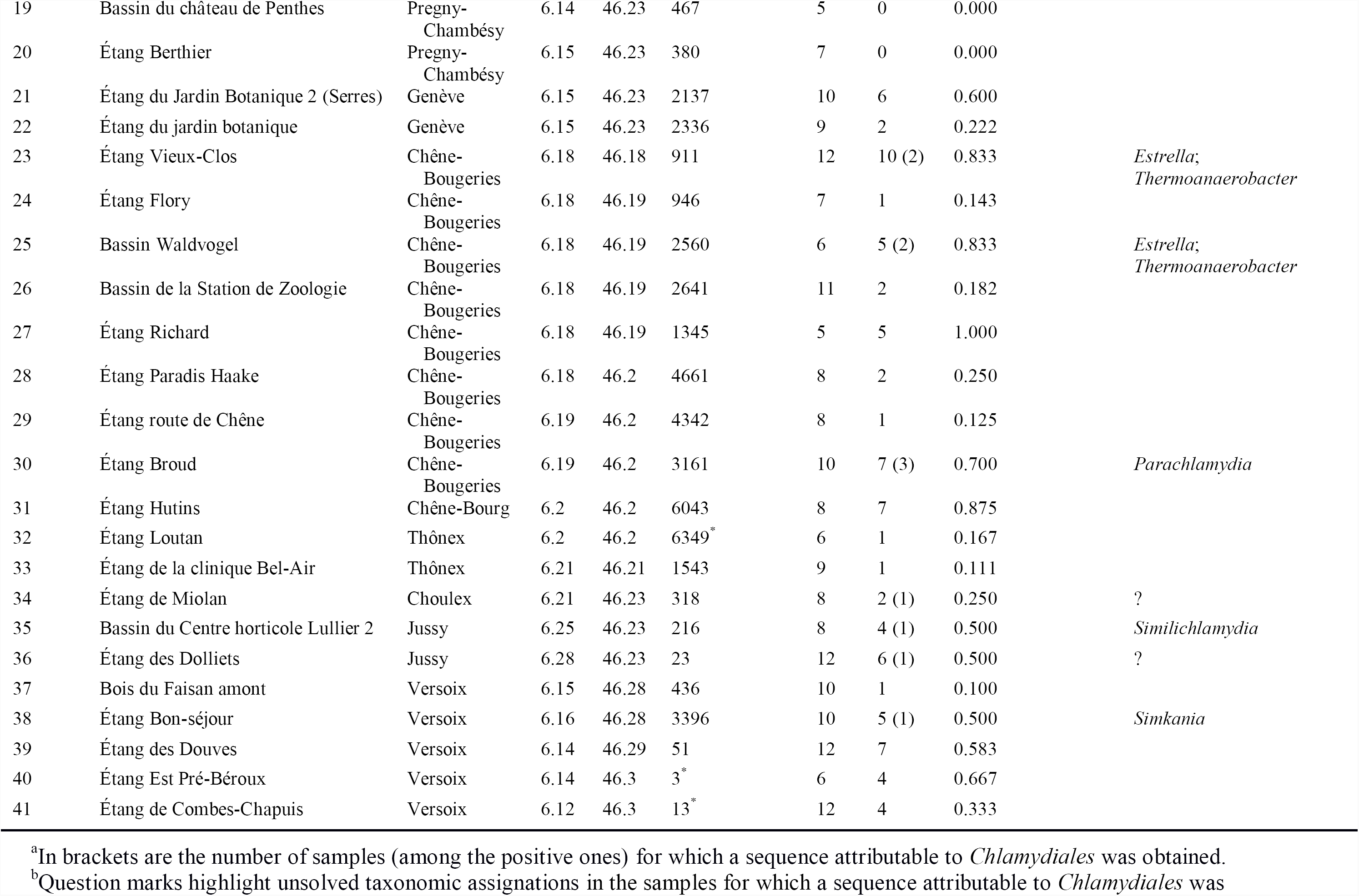
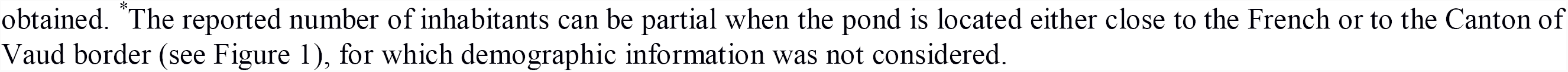
Information is reported for each pond including: experimental and actual name (Pond ID and Pond name, respectively), corresponding municipality, geographical coordinates, number of inhabitants (Nr. inh.s) within 1 km radius, number of *B. bufo* tadpoles sampled (N), number of CLOs-positive samples (P), observed infection rates (IR), and *Chlamydiales* taxonomic assignments at the genus-level.

#### Pan-Chlamydiales real-time PCR assay

A *Chlamydiales*-specific Real-Time PCR [44] was used to detect and amplify the DNA fragment from 207 to 215 bp belonging to the *Chlamydiales* 16S rRNA-encoding gene. Quantification was performed using a plasmidic 10-fold-diluted positive control tested in duplicate. Amplification reactions were performed in a final volume of 20 μl, containing: (i) iTaq Universal Probes Supermix with ROX (Bio-Rad, Reinach, Switzerland); (ii) 0.1 μM concentration of primers panCh16F2 (5′- CCGCCAACACTGGGACT-3′) (the underlined bases representing locked nucleic acids) and panCh16R2 (5′-GGAGTTAGCCGGTGCTTCTTTAC-3′) (Eurogentec, Seraing, Belgium); (iii) 0.1 μM concentration of probe panCh16S (5′-FAM [6-carboxyfluorescein]- CTACGGGAGGCTGCAGTCGAGAATC-BHQ1 [black hole quencher 1]-3′) (Eurogentec); (iv) molecular-biology-grade water (Five Prime, Hilden, Germany); (v) 5 μl of sample DNA. Amplification started with an initial step of activation and denaturation at 95°C for 3 min, followed by 40 cycles at 95°C, 67°C and 72°C, each lasting 15 s, and was performed in a StepOne Plus real-time PCR system (Applied Biosystems, Zug, Switzerland). Samples with a threshold cycle value (*C*_*T*_) <35 were finally sequenced, as this is the observed limit for amplicon sequencing (Aeby & Greub, unpublished).

#### DNA sequencing of the PCR-positive samples

According to the manufacturer’s instructions, amplicons from positive samples were purified using the MSB Spin PCRapace (STRATEC Molecular, Berlin, Germany). The sequencing PCR assay was performed using a BigDye Terminator v1.1 cycle sequencing kit (Applied Biosystems, Zug, Switzerland), and with specific inner primers panFseq (5′-CCAACACTGGGACTGAGA-3′) and panRseq (5′-GCCGGTGCTTCTTTAC-3′). Amplification was performed after an initial denaturation step at 96°C for 1 min followed by 25 cycles at 96°C for 10 s and 60°C for 4 min. Purification of the sequencing PCR products was done using the SigmaSpin Sequencing Reaction Clean-up (Sigma-Aldrich, Buchs, Switzerland), and the sequencing was performed in a 3130xL genetic analyzer (Applied Biosystems). Sequences were analysed and blasted using the Geneious software [45,46].

### Global spatial autocorrelation

Following taxonomic assignments, tadpoles’ infection rates (IRs) were computed for each pond. A global spatial autocorrelation analysis was then conducted to investigate the presence of clusters or dissimilarities (i.e. the existence of spatial groups with similar IRs values, or, on the contrary, the tendency of similar values to stay far away in space), under the null expectation of a random spatial distribution of infection. Moran Scatter plots [47,48] were then constructed testing different weighting criteria (in particular, using the mean IR from the first two, four, six, eight and ten nearest ponds, respectively), and Moran’s I [49] was estimated in each weighting scenario as the slope of the linear regression between weighted and observed IRs. Both observed and weighted IRs were centred prior to the analysis. Under the null hypothesis (*h*_*0*_) of a Moran’s I equal to zero and a significance threshold (α) set to 0.05, statistical significance of observed Moran’s I was derived by permuting IRs over the landscape for 9999 times, re-estimating Moran’s I in each permutation, deriving a Moran’s I reference distribution, and computing the pseudo *p*-value associated with the observed Moran’s I in each weighting scenario. Analysis was performed using a self-made script written in the R programming language [50,51].

### Human population density around ponds

The number of inhabitants residing within 1 km radius from each pond (surface: ~3.14 km^2^) was derived from the Federal Statistical Office database for the Republic and Canton of Geneva (www.bfs.admin.ch; see Table 1), and for the year 2013. To test for the existence of a spatial relationship between human and CLOs occurrence, the Pearson’s product moment correlation coefficient (r) was estimated between the number of inhabitants and the observed IRs, and a correlation test (*h*_*0*_: *r*=0; α=0.05) was performed through the function cor.test as implemented in the stats R package [50]. Ponds 1, 32, 40 and 42 were discarded from analysis given partial information about the number of inhabitants in the surrounding area.

### Group comparison analysis

Thirty-two categories describing land cover were derived from the territorial information system in Geneva (SITG) database (http://ge.ch/sitg/sitg_catalog/geodataid/1133) at a 10 m resolution. Proportions of each land cover category were then computed around the sampling sites as a function of a selected radius (i.e. buffer). In particular, buffers from 20 m up to 3 km (total: 299) were tested, and the R function extract [52] was used to extrapolate the land cover categories from the buffer circles.

To group ponds with similar characteristics, hierarchical clustering was performed on the resulting land cover proportions with the R function hclust [50], by relying on the “average”, “complete”, “single”, “Ward1” and “Ward2” clustering methods [53,54]. The Silhouette method [55] was used to identify the optimal number of clusters, as well as the ponds’ membership within the groups.

When two groups of ponds were identified, independent t-tests or Wilcoxon rank sum tests were run with the R functions t.test and wilcox.test, respectively; on the contrary, one-way ANOVAs or Kruskal-Wallis rank sum tests were performed with aov or kruskal.test [50] in the presence of more than two clusters. Test choice was driven by firstly checking IRs for normality and homoscedasticity. *P*-values from the analyses performed with the same cluster method were corrected for multiple testing with the “Benjamini-Hochberg” (BH) method through p.adjust [50]. Buffer scenarios with at least one group composed by a single pond were discarded given impossibility of assessing normality.

### Beta regression models

Association between land cover and IRs was also investigated through univariate beta regression models [56]. To ease interpretation of regression coefficients (*β*), previously obtained land cover proportions were aggregated into five classes prior to analysis (Supplementary Table 1): “Managed vegetation”, accounting for human-managed green areas; “Vegetation near water”, referring to species communities well-adapted to live into/or in close proximity with water surfaces; “Forests”, grouping forest species; “Urban environment”, encompassing highly urbanized areas; and “Open fields”, enclosing natural vegetation different from forests. Association tests (*h*_*0*_: *β*=0; α=0.05) were then performed separately for each class, and for each buffer used. The function betareg [56] was used to perform the tests, and observed IRs were transformed prior to the analyses to account for the presence of extreme values (zeros and ones) [57]. As before, *p*-values from the tests involving the same aggregated land cover class were corrected for multiple testing with the BH method.

## Results

We found 145 tadpoles (i.e. 38.7% of the samples) positive for the presence of *Chlamydiales* infection. Positive samples occurred across 36 sampling sites (i.e. 87.8% of the ponds sampled), with 5 ponds (12.2%) displaying no evidence of *Chlamydiales* occurrence, and infection rates spanning from 0 to 100% (Table 1 and Figure 2). Moran’s I analysis indicated the absence of significant spatial autocorrelation in the observed infection rates, regardless of the weighting scenarios used (Figure 3). In addition, no significant relationship was observed between human presence and observed IRs (*r*=–0.145; 95% confidence intervals: –0.448, 0.187; *t*=–0.869; df=35; *p*-value=0.391), with ponds displaying IR>0.5 being located in both scarcely and highly populated areas (see ponds 39 and 23 with ~50 and 900 inhabitants/3.14 km^2^, respectively, and ponds 31 and 17 with ~6000 and 7100 inhabitants/3.14 km^2^, respectively; Supplementary Figure 1).

**Figure 2.**
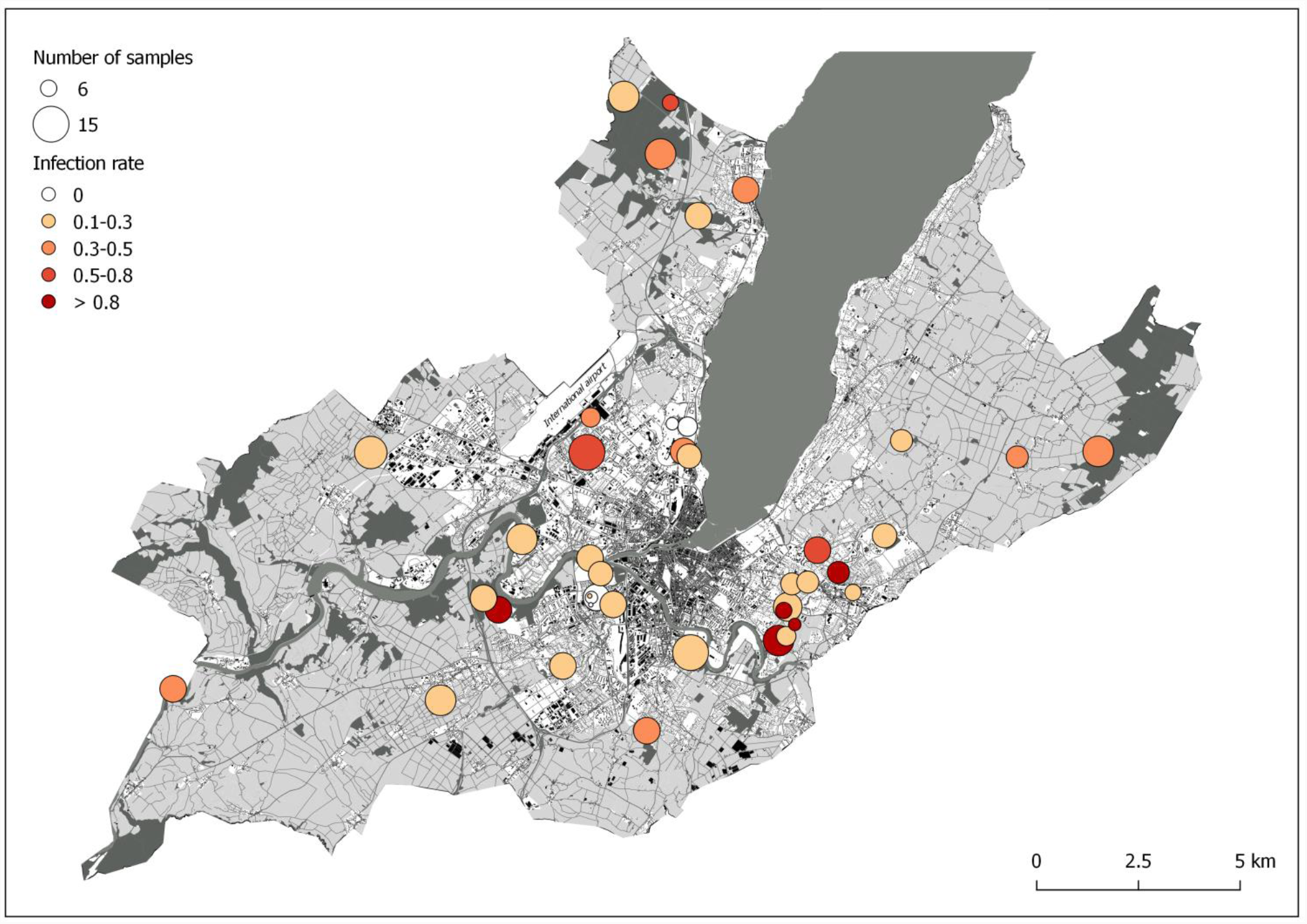
Observed infection rates over the Geneva metropolitan area. The circles represent sampling sites (see Figure 1), with size proportional to the number of tadpoles sampled, and hue intensity following the gradient in the observed infection rates.

**Figure 3.**
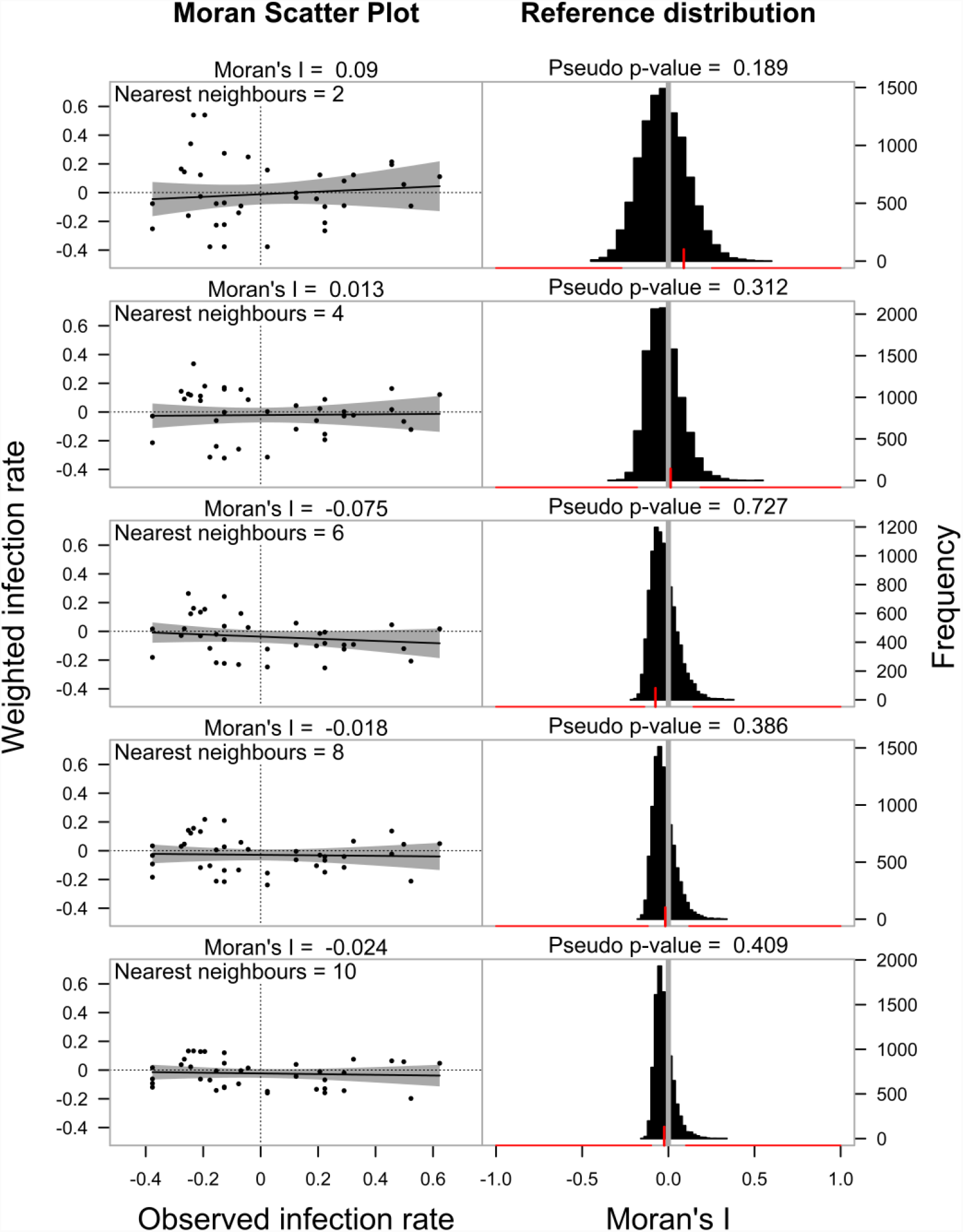
Global autocorrelation analysis results. Left column reports the Moran Scatter Plots obtained using the first two, four, six, eight and ten nearest neighbours (i.e. ponds), respectively. Right column reports the Moran’s I reference distributions as obtained for each weighting scenario by permutation tests. The red vertical tick highlights the position of the observed Moran’s I in the reference distribution; a grey vertical line is drawn to show I=0 (i.e. the null hypothesis). Red horizontal lines pinpoint percentiles 2.5 and 97.5 of the reference distributions, underlining the range of significant I values.

Only Wilcoxon and Kruskal-Wallis rank sum tests were performed on the groups of ponds defined with clustering analysis due to non-normality and/or heteroscedasticity. After multiple testing correction, none of the land cover-based group showed any significant difference in IRs (Figure 5). Likewise, none of the aggregated land cover variable showed a significant association with IRs after multiple testing correction in the beta regression analysis (Figure 6).

**Figure 4.**
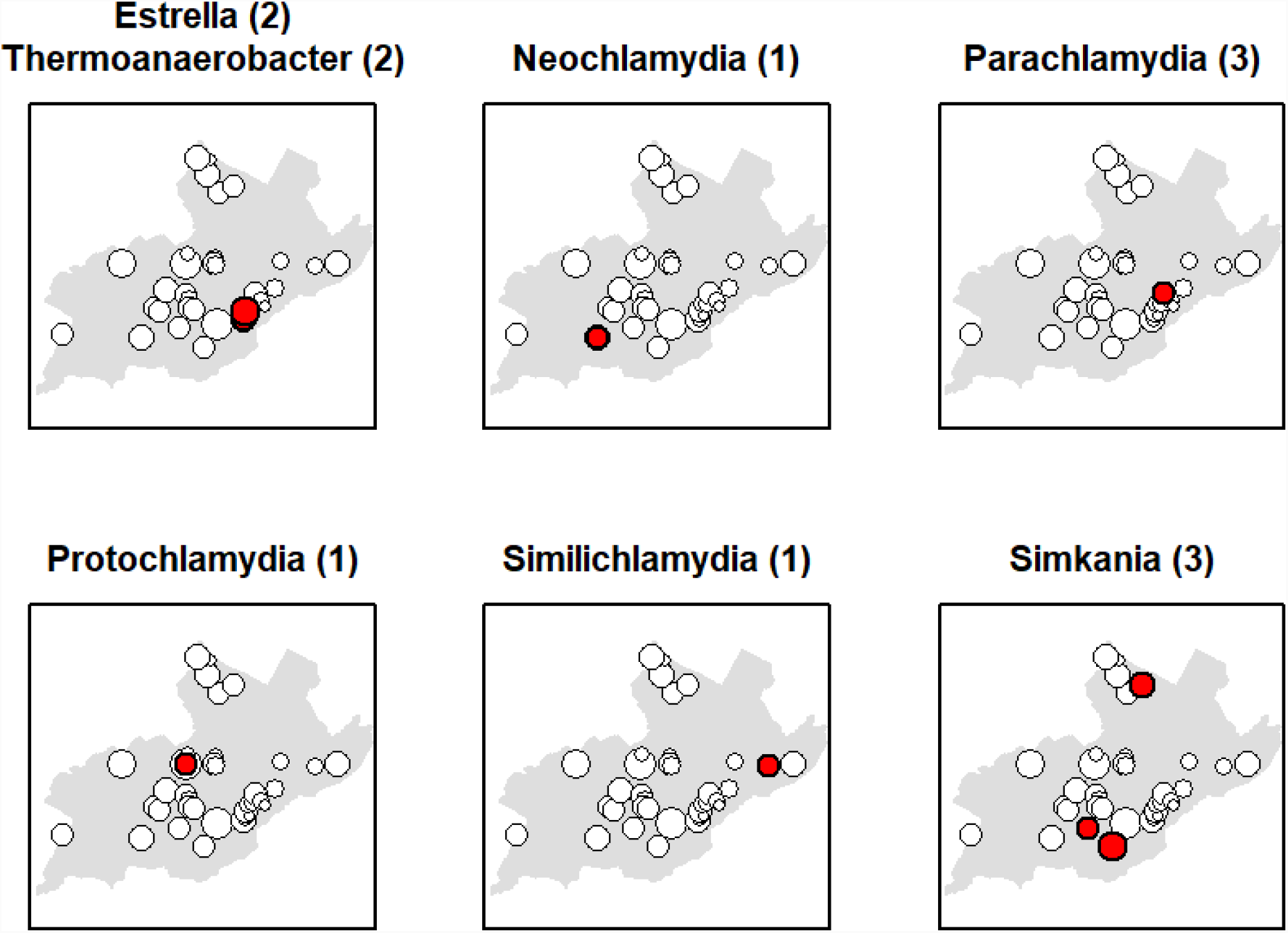
Spatial occurrence of the observed CLOs and *Thermoanaerobacteriaceae* genera is highlighted by red circles. The number of infected samples (i.e. tadpoles) is reported for each bacterial genera in brackets (see Table 1), and refers to the highlighted ponds (e.g. three tadpoles are positive for the genus *Parachlamydia* from the same highlighted pond). The grey area in the background represents the Geneva metropolitan area, and the size of the circles is proportional to the number of tadpoles sampled in each sampling site (see Figure 2).

**Figure 5.**
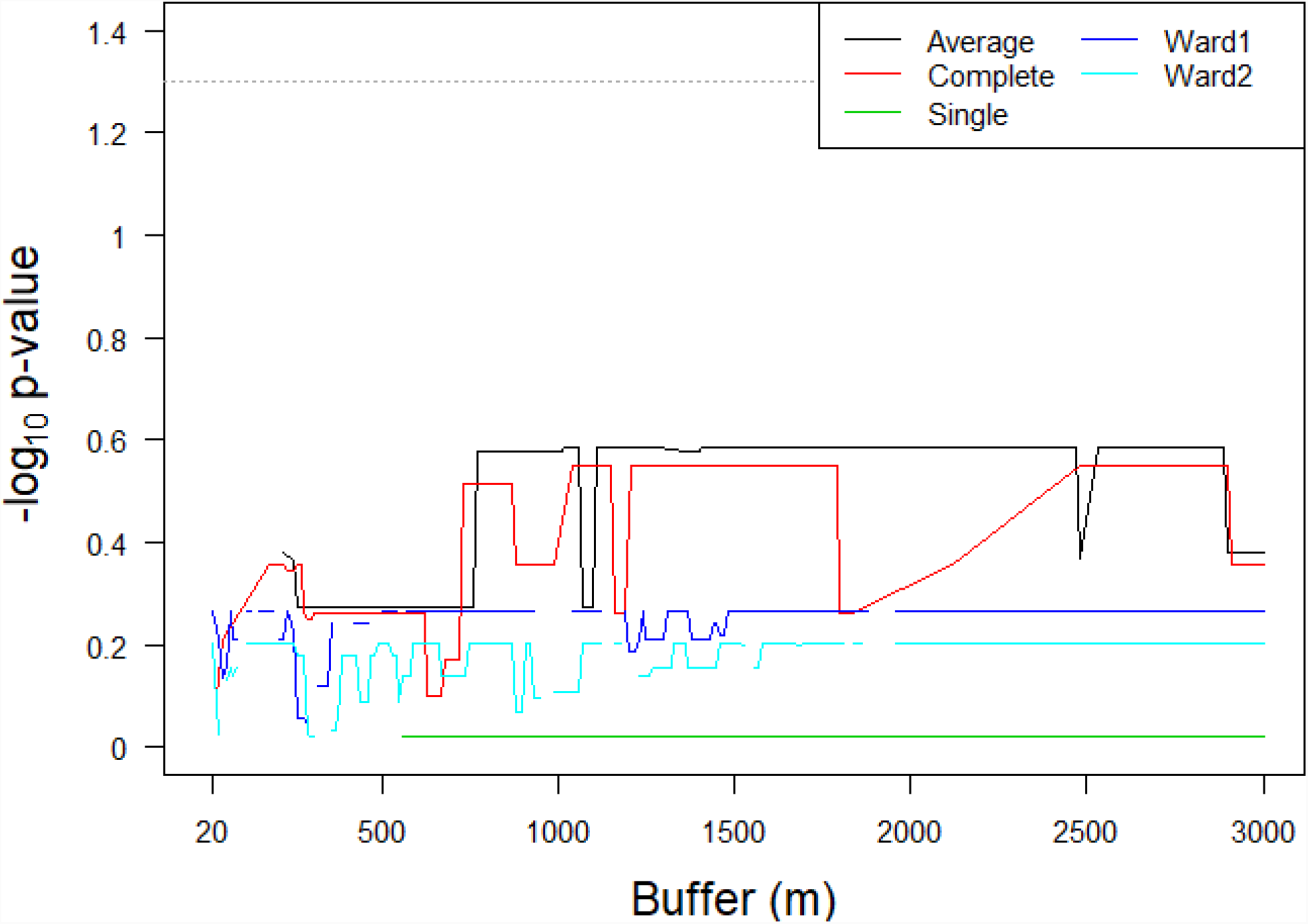
Results of the group comparison tests. *P*-values are reported on the logarithmic scale, after multiple testing correction, and as a function of both the buffer (i.e. radius) used for characterizing land cover around the sampling sites, and the clustering method used to classify ponds into environmental groups. In the uppermost part of the plot, the dotted line indicates the used significance threshold. Line discontinuities depict tests not run (i.e. where at least one group was constituted by a single pond; refer to main text for explanation).

**Figure 6.**
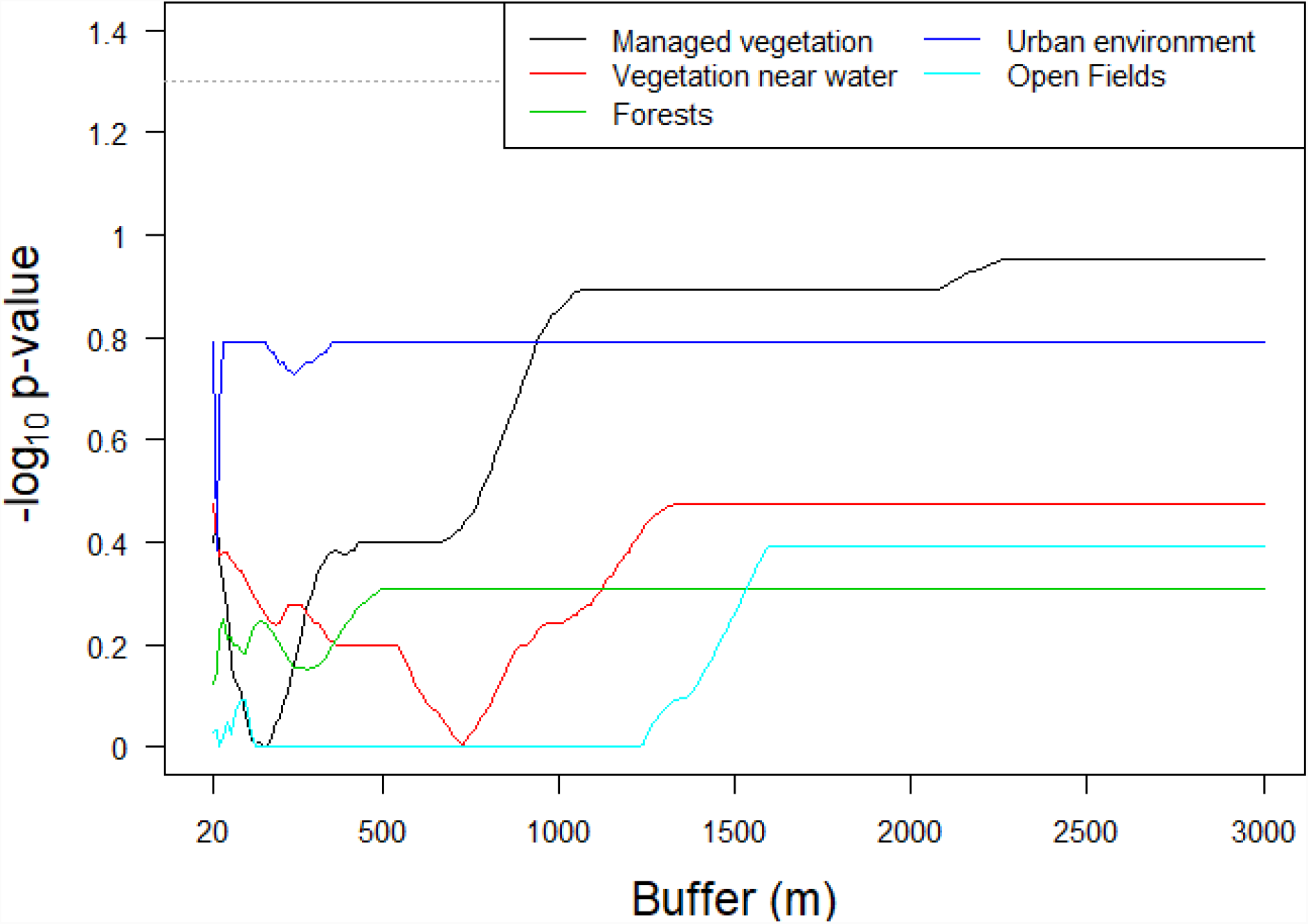
Results of the beta regression analysis. *P*-values associated with the estimated regression coefficients are reported on the logarithmic scale, after multiple testing correction, and as a function of both the buffer (i.e. radius) used for characterizing land cover around the sampling sites, and the aggregated land cover category. In the uppermost part of the plot, the dotted line indicates the used significance threshold.

Out of the 145 positive samples, 16 presented a *C*_*T*_ value <35, and were subsequently sequenced at the 16S ribosomal RNA gene for taxonomic identification. Taxonomic attribution was possible at a family-level lineage for 13 samples. In particular, six were found to be positive for *Parachlamydiaceae*, three for *Simkaniaceae* and two for *Criblamydiaceae*. The remaining two samples were found positive for the family *Thermoanaerobacteriaceae*, genus *Thermoanaerobacter*, which does not belong to *Chlamydiales*. Among the six samples infected by *Parachlamydiaceae*, one was positive for the genus *Similichlamydia* (as retrieved from Pond 35), one for the genus *Neochlamydia* (from pond 3), one for the genus *Protochlamydia* (from Pond 17), and three for the genus *Parachlamydia* (as observed in pond 30, with two sequences being highly similar with less than 1% divergence in the 16S rRNA sequence). All the samples infected by *Simkaniaceae* were assigned to the genus *Simkania* (from ponds 7, 15 and 38, respectively). Finally, samples infected by *Criblamydiaceae* and positive for *Thermoanaerobacteriaceae* were assigned to the genera *Estrella* and *Thermoanaerobacter*, respectively, and retrieved from Ponds 23 and 25 (Table 1 and Figure 4). Due to low sequencing quality, no lineage could be identified for the samples coming from Ponds 1, 34 and 36.

## Discussion

The order *Chlamydiales* comprises bacterial agents of important human and animal diseases, as well as emerging pathogens which affect a broad spectrum of hosts [1,3,26]. To our knowledge, the present study reports the first observation of CLOs infection in tadpoles’ populations of the common toad species *B. bufo*.

Notably, among the operational taxonomic units found are CLOs assigned to the genus *Parachlamydia*, which characterized the microbiome of a Pyrenean midwife toad population unable to recover from an infection by the highly aggressive fungus *Batrachochytrium dendrobatidis* [25]. Considering co-occurrence of Chlamydiae and *B. dendrobatidis* was also observed in a *X. tropicalis* population undergoing epizootic disease dynamics [12], an association was proposed between the skin microbiome of amphibians and *B. dendrobatidis* infection outcome [25]. Given the emerging role of *B. dendrobatidis* in the current global amphibian biodiversity crisis [58], *Parachlamydia* occurrence might pinpoint a potential vulnerability for the *B. bufo* populations under study which should deserve attention for conservation.

The genera *Simkania* and *Neochlamydia* encompasses recognized emerging pathogens for both humans and animals. Particularly, *Simkania* species were associated with respiratory deficit in humans and epitheliocystis in fish, and *Neochlamydia* species with ocular diseases in domestic cat and epitheliocystis [3]. So far, there is weak evidence about *Estrella* involvement as a human pathogen, even if this recently discovered bacterial genus is still understudied [59]. Such findings would suggest a potential role for *B. bufo* as a host reservoir for CLOs, and an ad hoc monitoring program might be beneficial for both biodiversity conservation and public health purposes. Nevertheless, no evident relationship seems to exist between observed infection rates and population density in the study area, even if further studies would be advisable relying on a bigger sample size to obtain more robust evidences in this regard; furthermore, particular consideration should be accorded to *Le Marais* and *Étang Hutins* (Pond 17 and Pond 31, respectively) given their combination of high IRs and number of inhabitants (Supplementary Figure 1).

Several studies support the paradigm of microbial biogeography “everything is everywhere, but, the environment selects” for *Chlamydiales* [1,3,42]. Coherently with such literature statements, the apparent absence of a clear spatial pattern, together with the rather ubiquitous occurrence of infection in *B. bufo*, would suggest a cosmopolitan distribution for the vectors and their endosymbionts even at a local geographical scale (see Figures 2 and 3), thus comforting the expectation “everything is everywhere”. Nevertheless, our attempt to investigate how “the environment selects” was more tricky, and no association was found with land cover typologies able to explain the observed CLOs distribution (see Figures 5 and 6). At this regard, we believe further studies should focus on the influence of local environmental conditions on *Chlamydiales* occurrence, especially tacking into consideration alternative variables with proved effects on amoebae distributions (e.g. precipitation, temperature, moisture, pH and dissolved oxygen) [36–40]. Ideally, the description of CLOs-specific niches would provide fine-scale predictive distribution maps of *Chlamydiales* and their vectors, with straightforward applications in both public health preventive strategies, and prioritization of susceptible animal host populations for conservation.

To conclude, the present work candidates the amphibian species *B. bufo* as a new host for CLOs, provides a first estimate of CLOs infection rate in *B. bufo* tadpole populations from a urban environment, agrees with literature findings concerned with *Chlamydiales* ubiquitous distribution, while apparently excluding the land cover as a selective variable for *Chlamydiales* occurrence in the studied area.

## Acknowledgements

We warmly thank the residents of the city of Geneva who allowed us to access their private pond for sampling. The study was funded by the GELBERT Foundation in Geneva, Switzerland (URBANGENE project no 088-2013), by the Grand Genève cross-border metropolitan area, and by the Direction générale Agriculture & Nature (DGAN) du Département de l’Environnement, des Transports et de l’Agriculture (DETA) of the State of Geneva, Switzerland.

